# mTORC2 mediate FLCN-induced HIF2α nuclear import and proliferation of clear cell renal cell carcinoma

**DOI:** 10.1101/2020.01.13.905521

**Authors:** Xuyang Zhao, Yadong Ma, Jie Cui, Haiyang Zhao, Lei Liu, Yueyuan Wang, Pengxiang Min, Lin Zhang, Yongchang Chen, Jun Du, Yujie Zhang, Luo Gu

## Abstract

Clear cell renal cell carcinoma (ccRCC), as the most important type of renal carcinoma, has a high incidence and easy metastasis. Folliculin (FLCN) was identified as a tumor suppressor gene. Its deletions and mutations are associated with a potential risk of kidney cancer. At present, the specific molecular mechanism of FLCN-induced proliferation, invasion and migration in clear cell renal cell carcinoma remains elusive.

In this study, we demonstrated that FLCN controled cell proliferation, invasion and migration through PI3K/mTORC2 pathway. FLCN combined with HIF2α in various normal and cancerous renal cells, and mTORC2 mediate FLCN effectively alleviated the deterioration of renal cancer cells by degrading HIF2α. Silencing of FLCN showed promotion of HIF2α protein expression, which in turn led to an increase in downstream target genes *Cyclin D1* and *MMP9*. Moreover, when interfering with siFLCN, HIF2α degradation rate was delayed, and the time of entry into the nucleus was advanced. Taken together, our study illustrated that mTORC2 promoted the specific molecular mechanism of HIF2α by down-regulated FLCN, and might be a new therapeutic target against renal cancer progression.

## 1. Introduction

Renal cell carcinoma is a malignant tumor originating in the renal tubular epithelial system, and most of which are clear cell renal cell carcinoma (~75%) ^1^. Cell viability is important for various physiological processes such as embryonic development, angiogenesis, and tumor proliferation, invasion and metastasis. Birt-Hogg-Dubé syndrome is caused by inactivating mutations of FLCN, a tumor suppressor gene which encodes folliculin ^2^. Common causes of Birt-Hogg-Dubé syndrome are lung cyst, spontaneous pneumothorax, skin fibrofolliculomas and renal cancer ^3^. It is also reported that inactivation of FLCN is an initial step in the development of renal tumors in BHD ^4^. Work by Sok Kean Khoo and Laura S. Schmidt have confirmed a tumor suppressor role for FLCN ^5–7^. Similarly the FLCN-deficient animal models ^8^ showed activated mTOR and its downstream pathway effectors ^9^. These paper suggested that FLCN-deficient kidney tumors showed activation of mTOR and AKT ^10^. And this regulation mechanism is also established in humans ^2,11,12^.

Hypoxia-inducible factor (HIF) is a crucial mediator of hypoxic adaptation. Previous studies have shown that renal tumor-associated gene VHL plays a decisive role in regulating HIF expression ^13,14^. Similarly, research data also shows that there are many links between FLCN and HIF1α ^8,15^. Our pretest result indicated that the *FLCN* gene acts as a regulator of renal tumors and its knockdown resulted in a significant increase in HIF2α protein levels. *Cyclin D1* and *MMP9* have been identified as HIF2α-regulated genes which can mediate cancer cell proliferation and invasion ^16–18^.

In this study, we hypothesized that FLCN may regulate HIF2α through binding to HIF2α to suppress expression of Cyclin D1 and MMP9 during cell proliferation and invasion. Here, we showed that FLCN regulates the nuclear export timing of the transcription factor HIF2α by forming a complex with HIF2α. Knockdown of *FLCN* caused the HIF2α to enter the nucleus in advance, further exacerbated the aggressive behavior of the tumor. In addition, FLCN/HIF2α was also regulated by PI3K/mTORC2 signaling pathway. These findings revealed a novel relationship between FLCN and HIF2α in clear cell renal cell carcinoma proliferation, invasion and metastasis.

## 2. Materials and methods

### 2.1 Ethics statement

All immunohistochemistry assays with human tumour specimens were conducted under the institutional guidelines of Jiangsu Province.

### 2.2 Cell lines and cell culture

Normal renal tubular epithelial cell line HK-2 and human clear cell renal cell carcinoma lines 786-O and ACHN were obtained from the Cell Biology Institute of Chinese Academy of Sciences (Shanghai, China).

HK-2, and ACHN cells were cultured in DMEM (Hyclone, ThermoScientific, Waltham, MA) supplemented with 10% fetal bovine serum (FBS; Gibco, Carlsbad, CA, USA) at 37 °C in 5% CO_2_. 786-O cells were cultured in RPMI 1640 (Hyclone, ThermoScientific, Waltham, MA) supplemented with 10% fetal bovine serum at 37 °C in 5% CO_2_. The cells were grown on 10 cm dishes for subsequent experiments.

### 2.3 Plasmids and siRNAs

The full-length Epas1 plasmid was purchased from Youbio (Hunan, China). The pCMV-FLAG-FLCN plasmid was constructed as previously reported, using the following primer set. The above constructs were confirmed via DNA sequencing. The cells were seeded in six-well plate, cultured to 80%~90% confluence, and then transfected with Epas1 and pCMV-FLAG-FLCN plasmids respectively by using FuGENE HD Transfection Reagent (Promega Corporation, Madison, WI, USA).

Non-specific control siRNA and siRNAs for FLCN and EPAS1 were purchased from GenePharma (Shanghai, China). The cells were transfected with siRNA Duplex oligonucleotides using Lipofectamine 2000 (Invitrogen), according to the transfection method provided by the manufacturer. Following siRNAs were listed in TABLE 1.

**TABLE 1.**
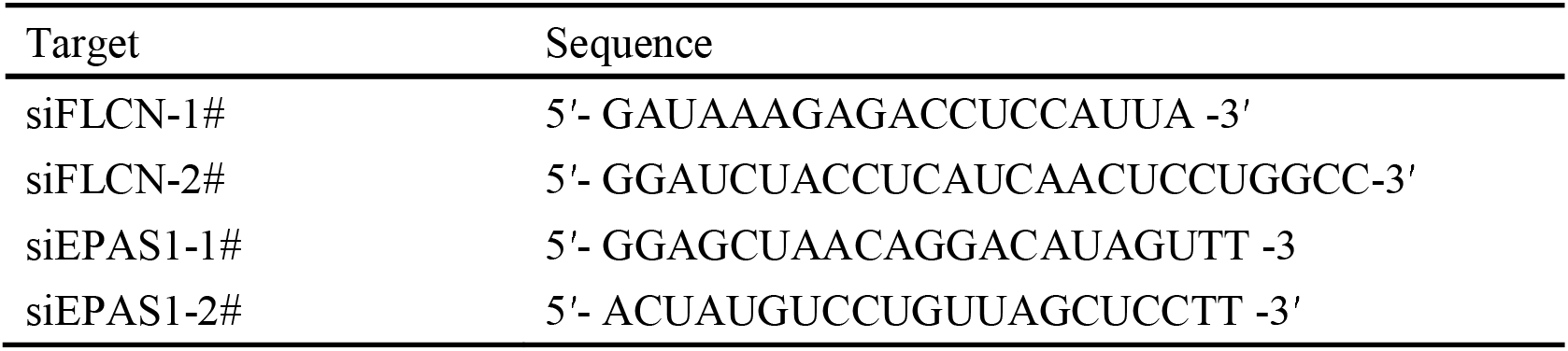
siRNAs sequences used for transfection.

### 2.4 Reagents and antibodies

Cyclohexamide (CHX) was purchased from Sigma (USA), MG-132, MK2206 (AKT inhibitor), KU0063794 (mTORC1 and mTORC2 inhibitor), Rapamycin (mTORC1 inhibitor), LY294002 (PI3K inhibitor) were purchased from ApexBio. Rabbit monoclonal antibodies against AKT, phospho-AKT (Thr308, Ser473), Histone 3, MMP9 and HIF2α were purchased from Cell Signaling Technology. Mouse monoclonal antibody against HIF2α was purchased from Millipore for immunofluorescence. FLCN rabbit pAb was purchased from Cell Signaling Technology for western and ABcam for immunoprecipitation. GAPDH rabbit mAb and β-actin mouse mAb antibodies were purchased from Santa Cruz Biotechnology. HRP conjugated secondary antibodies were purchased from Jeckson Immunoresearch Laboratories.

### 2.5 Immunoprecipitation (IP) and western blotting

For Western blotting, equal amounts of protein were run on SDS polyacrylamide gels and transferred to nitrocellulose membrane. The resulting blots were blocked with 5% non-fat milk (in TBST) and incubated with primary antibodies overnight at 4◻. Then protein bands were detected by incubating with secondary antibodies for 1-2h at room-temperature and visualized with ECL reagent (Millipore, Billerica, MA, USA) by Chemi Doc XRS+gel imaging system (Bio-Rad, USA). Densitometry analysis was performed using Quantity One software, and band intensities were normalized to those of β-Actin and GAPDH.

IP was performed as previously described^19^. Briefly, the cell lysates were centrifuged to remove the cell debris and then were incubated with beads (Abmart) for 1-2 h. Endogenous FLCN and HIF2α was immunoprecipitated using an anti-FLCN or anti-HIF2α polyclonal antibody. The beads were boiled after extensive washing; resolved via SDS-PAGE gel electrophoreses, and analyzed via immunoblotting. The protein concentration was detected using the Odyssey system.

### 2.6 Cytoplasmic and nuclear protein extraction

Cytoplasmic and nuclear proteins were obtained using the Nuclear and Cytoplasmic Protein Extraction Kit (Beyotime) according to the manufacturer’s instructions. Cells were harvested by centrifugation and resuspended in cytoplasmic extraction agent A. The solution was vigorously vortexed and then incubated on ice for 10 minutes. Then, cytoplasmic extraction agent B was added to the cell pellet. The pellet was vortexed and incubated on ice for 1 minute. The pellet was vortexed again and centrifuged for 5 minutes at 5000 rpm. The Supernatant was collected as cytoplasmic extract. The insoluble (pellet) fraction was suspended by nuclear extraction agent. After vortexed several times, the mixture was centrifuged for 10 minutes at 12000 rpm. The supernatant was collected as nuclear extract.

### 2.7 Real◻time quantitative PCR

Total RNA was extracted using Trizol reagent (Invitrogen) and reversely transcribed with HiScript^®^Q RT SuperMix for qPCR (Vazyme, Nanjing, China) according to the protocol. Real◻time PCR analyses were performed with AceQ^®^ qPCR SYBR^®^ Green Master Mix (High ROX Premixed) (Vazyme) on ABI StepOne™ Real◻Time PCR System (Applied Biosystems, Foster City, CA) at the recommended thermal cycling settings: one initial cycle at 95°C for 10 minutes followed by 40 cycles of 15 seconds at 95°C and 60 seconds at 60°C. The gene expression levels were calculated with Rt (2^−ΔΔCT^) values by StepOne Software v2.1 (Applied Biosystems). Primer sequences used in qRT◻ PCR were listed in TABLE 2.

**TABLE 2.**
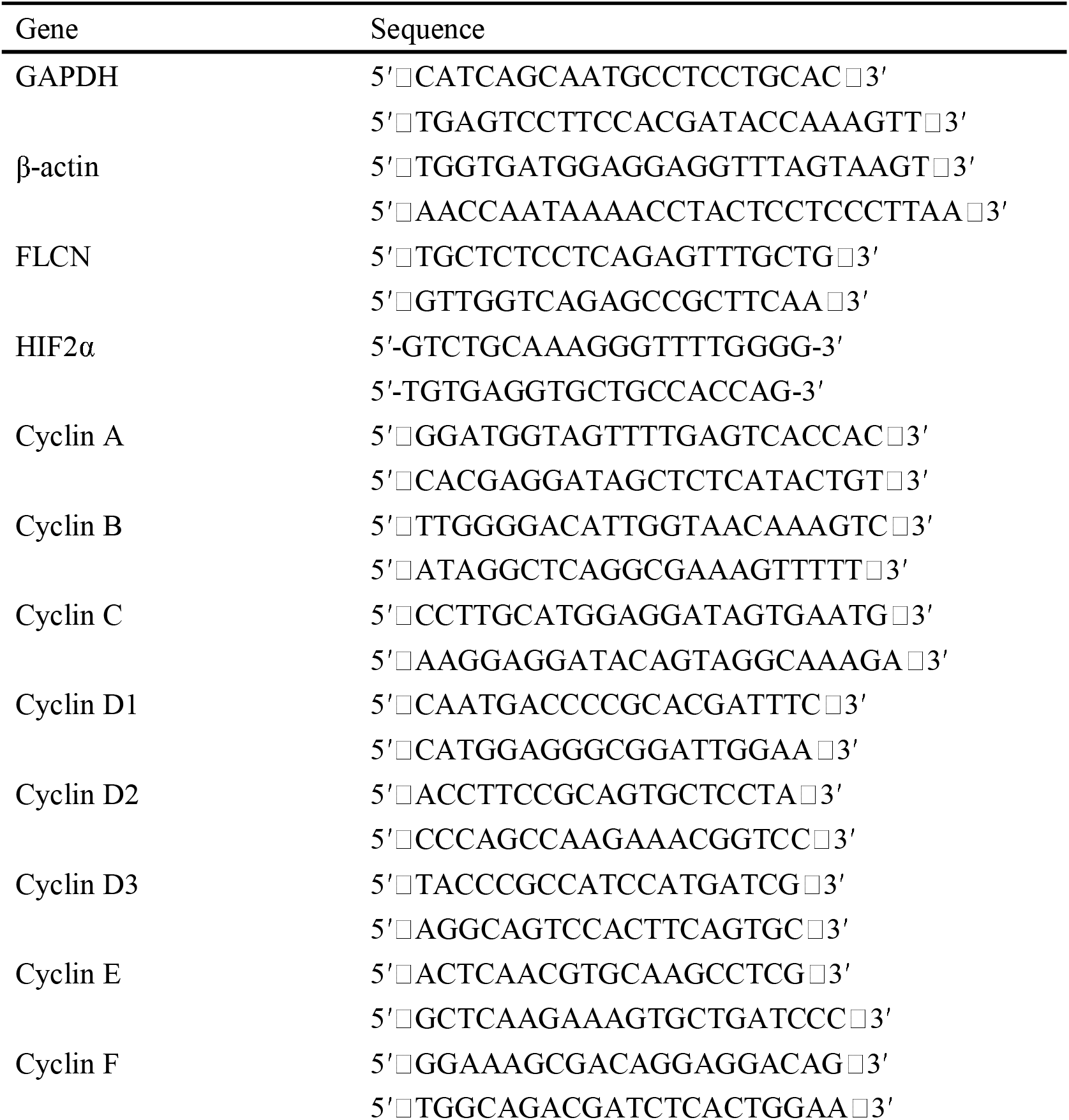
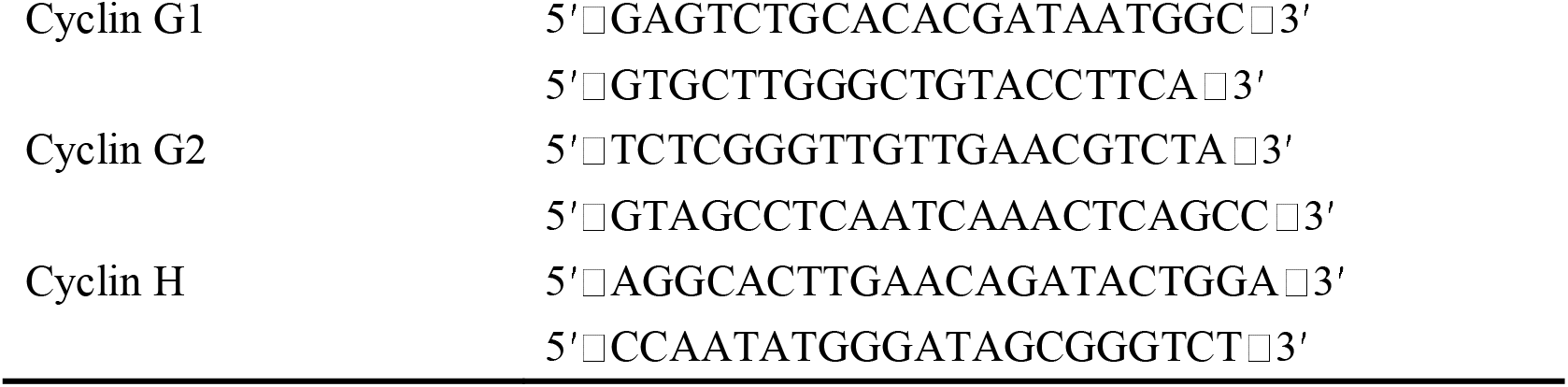
Primer sequences used for qRT-PCR.

### 2.8 Immunofluorescence microscopy

For immunofluorescent staining, cells were seeded on coverslips. The cells were transfected with or without siFLCN and FLCN plasmids for the indicated time, fixed in ice-cold 4% paraformaldehyde for 20 min at room temperature, rinsed with 3×phosphate-buffered saline (PBS) for 5 min and permeabilized with 0.1% Triton X-100 before blocking in 1% BSA for 1 h at room temperature. The cells were incubated with primary antibodies at 4 °C overnight, washed, and then incubated with Alexa or FITC-coupled secondary antibodies for 1.5 h at room temperature in a moist chamber. After washing with PBS, the samples were mounted with DAPI Fluoromount G (Southern Biotech, Birmingham, AL). Images were acquired using an Olympus BX51 microscope coupled with an Olympus DP70 digital camera and Carl ZEISS MicroImaging GmbH (Jena, Germany, 700). Images were representatives of three independent experiments.

### 2.9 Migration and Matrigel invasion assays

For wound healing assay, 786-O or ACHN cells were grown as described above and were plated in 6 well plates on glass cover slips. Approximately 24 h later, when the confluence reached 95◻~◻100 %, the cells were incubated overnight in DMEM and the wounding was performed by scraping through the cell monolayer with a 10 μL pipette tip. The Medium and the nonadherent cells were removed, and the cells were washed twice with PBS. Images were collected of the 0 h time point using an inverted microscope (Carl Zeiss Meditec, Jena, Germany). The Cells were permitted to migrate into the area of clearing for 6h, 12h and 24h in incubator. And then, the cells were removed from the incubator and imaged using the same inverted microscope. Care was taken to align the scratch along the y axis of the camera to aid subsequent image quantification.

For Matrigel invasion assay, the cell invasion assay was performed using Transwells (8 μm pore size, millipore) with inserts coated with Matrigel (50 mg/ mL, BD Biosciences). 786-O and ACHN cells (1.0 × 10^5^ cells/well) were seeded in the upper chambers with 0.1 mL matrigel and allowed to invade through matrigel for 12 h and 24 h. The cells remained on the membranes were fixed with 4% paraformaldehyde and stained with 0.5% crystal violet.

The cell pictures were taken by Nikon TS100 (Tokyo, Japan) and counted by Image J software. All assays were performed at least three times.

### 2.10 CCK-8 assay and Flow cytometry analysis

The Cells transfected with plasmids or siRNA were seeded at a density of 3.5 × 10^3^ cells per well into 96-well plate. After culture, the cells were washed, add 10 μl Cell Counting Kit-8 (CCK-8) per 100 μl culture medium and the plate was incubated in the dark for 2 h, followed by measurement of absorbance value at 450 nm using a microplate absorbance reader (Bio-Tek, Elx800, USA). The fold growth was calculated as the absorbance of drug treated sample/control sample absorbance ×100%.

Cell cycle analysis was performed by flow cytometry. Briefly, cells were harvested and fixed in 80% ice◻cold ethanol overnight. Then the cells were incubated with RNase A and propidium iodide staining solution at 37°C for 30 minutes in darkness. Subsequently, the stained cells were analysed using flow cytometry.

### 2.11 Immunohistochemistry

Renal cancer tissue microarrays were purchased from Outdo biotech (Shanghai, China). 75 cases of Renal clear cell carcinoma samples and their corresponding paracancerous tissue samples were used for immunohistological staining in our study. Briefly, after microwave antigen retrieval, microarray tissues were incubated FLCN (Proteintech), or HIF2α antibody (Sigma) overnight at 4°C, followed by 1 hour incubation with secondary antibody. The sections were developed in DAB solution under microscopic observation and counter stained with haematoxylin. Immunohistochemical staining results were taken by using Axioskop 2 plus microscope (Carl Zeiss). FLCN and HIF2α immunostaining was scored by immune reactive score (IRS) as described previously^20^.

### 2.12 Statistical analysis

Student’s *t*-test and repeated-measures were used to analyze differences between groups by using the SPSS statistical software program (Version 19.0; SPSS, Chicago, IL, USA). Data were presented as mean ± S.E.M. Values of *P* < 0.05 were considered statistically significant. All experiments were repeated at least three times.

## 3. Results

### 3.1 FLCN is involved in proliferation of ccRCC cells

Firstly, we tested whether FLCN plays a crucial role in the HIF2α expression in ccRCC cells and verified the definite mechanisms involved. We first detected the protein levels of FLCN in normal renal tubular epithelial cells and clear cell renal cell carcinoma by Western blotting. The results showed that FLCN was poorly expressed in the 786-O and ACHN ccRCC cells compared to renal tubular epithelial cells HK-2. However, HIF2α protein expression level was opposite to FLCN trends (Fig. 1A). The expression of the FLCN and HIF2α had been examined by Realtime-PCR analysis in three cell lines (Fig. 1B). Western blotting confirmed siRNA mediated specific knockdown of FLCN in 786-O and ACHN cells (Fig. 1C), and the results of CCK-8 assay showed that the knockdown efficiently promoted the cell viability (Fig. 1D).

**Figure 1.**
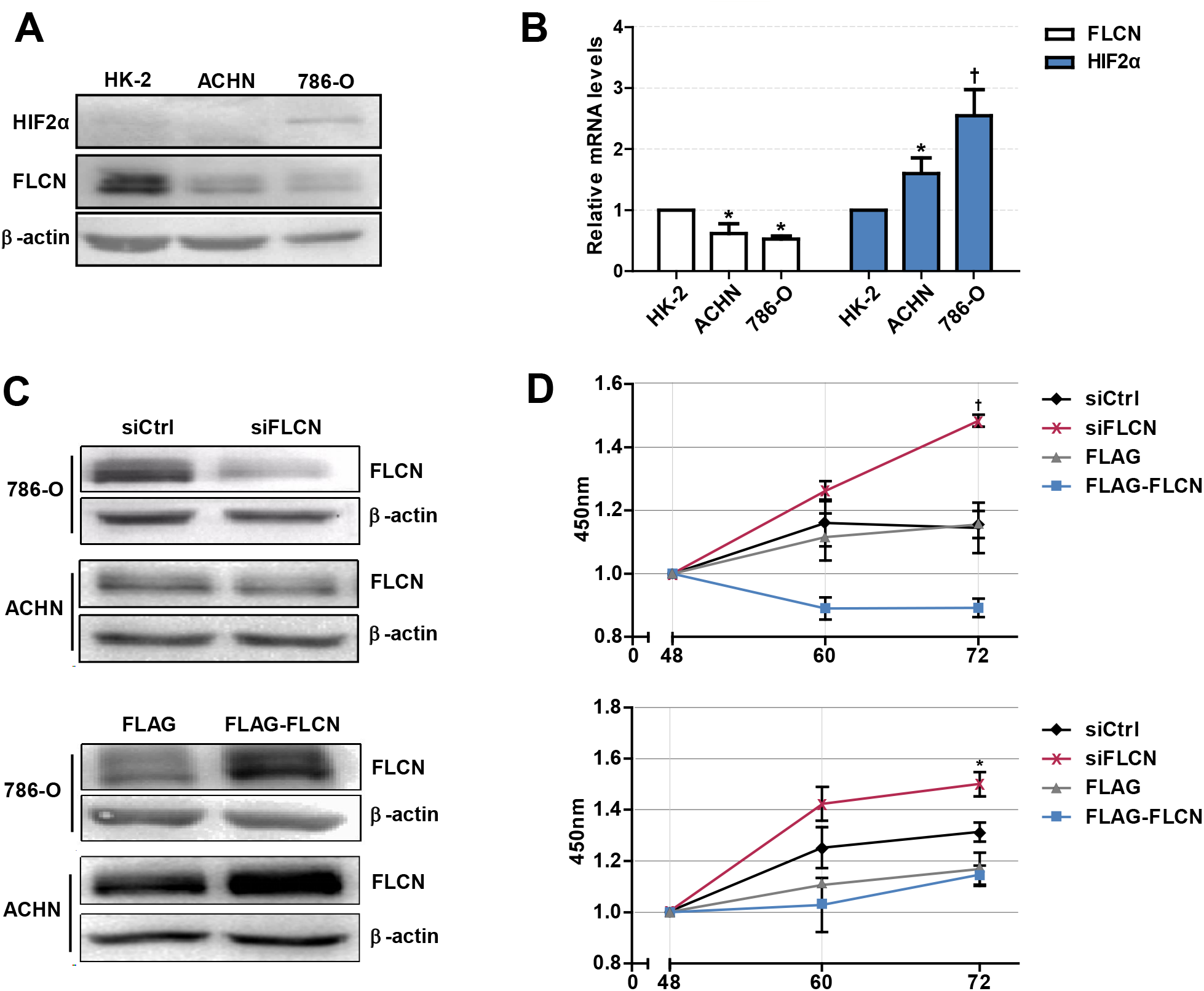
FLCN knockdown accelerates the proliferation of ccRCC cells. (A) Western blot detection of FLCN and HIF2α protein expression in ccRCC cell lines. (B) RT-qPCR analysis of FLCN and HIF2α mRNA expression in ccRCC cell lines. ccRCC cell lines including 786-O and ACHN were used in this experiment. Normal renal tubular epithelium cell line HK-2 was used as control. 786-O and ACHN cells were transfected with FLCN siRNA or overexpression vector for 48 h, and (C) the protein expression of FLCN was confirmed by Western blot. (D) Cell proliferation was assessed by CCK-8 assay. The data was from three independently repeated experiments. *: *P*<0.05, †: *P*<0.01.

### 3.2 FLCN negatively regulates ccRCC cell cycle and invasion in vitro

Since FLCN knockdown increased the viability of ccRCC cells, the second set of analyses examined the role of FLCN in cell motility. The results showed that FLCN knockdown affected the cell cycle in 786-O cells, and the number of cells in S phase was significantly increased (Fig. 2A left). In contrast, after transfection with the FLCN plasmid, the proportion of cells in S phase was reduced (Fig.2A right). The above experiment was also performed with ACHN cells, and similar results were obtained (Fig.2B). The results of wound healing assays showed that the rate of migration of cells transfected with siFLCN #1 and #2 was increased when it was compared to the control group in 786-O cells (Fig. 2C). FLCN overexpression gave the opposite results. The same conclusion was also drawn in ACHN cells (Fig. 2D). These results suggested a positive role for FLCN in regulating ccRCC cells migration, and compared with long-term migration rate (24h), the short-term (12h) rate was more obvious. We checked cell invasion by transwell assay, and found that FLCN knockdown increased the invasion as compared with control in 786-O and ACHN cells (Fig.2E).

**Figure 2.**
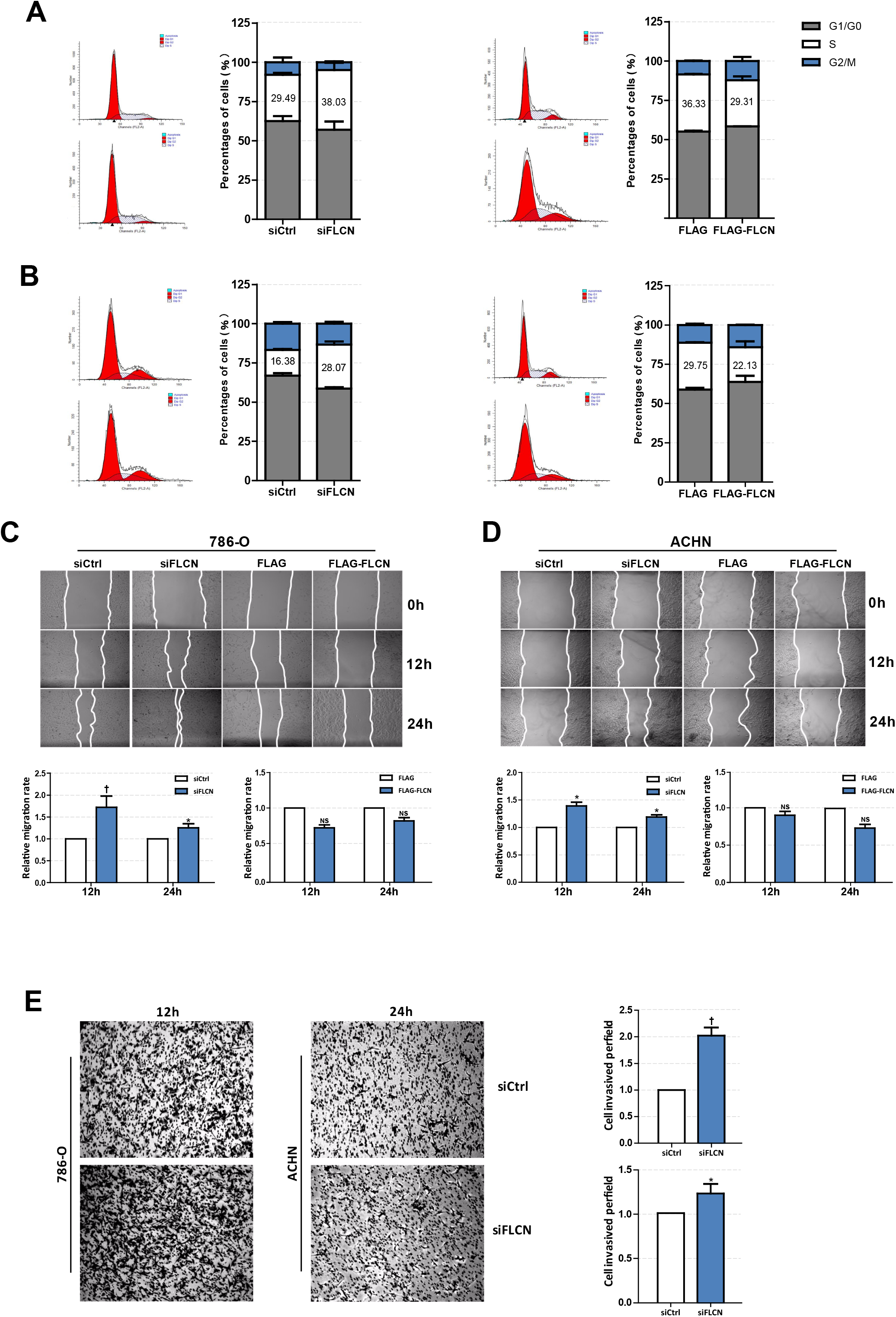
FLCN regulates ccRCC cell cycle and invasion *in vitro*. (A,B) Effects of FLCN on 786-O (A) and ACHN (B) cell cycle distribution. 786-O and ACHN cells were transfected with FLCN siRNA or overexpression vector for 48 h, flow cytometry assay was performed to test cell cycle. (C,D) A representative of wound healing assays in 786-O (C) and ACHN (D) cells transfected FLCN siRNA or overexpression vector for 6h or 12h (12h or 24h), and the quantification of cell migration rate was performed. (E)786-O and ACHN cells transfected with control siRNA or siFLCN for 12 or 24h, and the quantification of cell invasion rate was performed. The data was from three independently repeated experiments. *: *P*<0.05, †: *P*<0.01.

### 3.3 Cells lacking FLCN have elevated levels of HIF2α expression

To explore the mechanism of FLCN regulating cell proliferation and cell cycle, we focused on its putative interacting protein HIF2α. The silencing FLCN expression in 786-O cells with siRNA targeting FLCN, or the upregulating with Flag-FLCN plasmid was performed. The knockdown or overexpression efficiency was determined by Western blotting. The results showed that HIF2α and FLCN protein expression are negatively correlated (Fig. 3A). Conclusions in the ACHN cells are consistent with the above (Fig. 3B). In order to confirm HIF2α regulation by FLCN, the cells were treated with siFLCN or FLCN-overexpression plasmids, and then analysed for FLCN and HIF2α mRNA level by real-time PCR. The results showed that the expression of *FLCN* gene was obviously changed with the treatment, but none of this program caused a significant effect on HIF2α gene as compared with control (Fig. 3C and 3D). Then the expression of the genes of cyclin family (FLCN/HIF2α downstream cell-viability related genes) in HK-2 (normal), 786-O and ACHN (ccRCC) cells was detected. Consistent with expectations, the expression level of cyclin D1 was significantly higher than others (Fig. 3E). According to this, the follow-up experiment used cyclin D1 as a marker, to detect if HIF2α was directly downstream of FLCN. We co-transfected siFLCN/siHIF2α in 786-O cells, and detected *cyclin D1* gene expression levels by real-time PCR (Fig. 3F left). Similarly, overexpression of FLCN reduced *cyclin D1* gene expression levels, and under these conditions, co-transfections with Epas1 plasmid rescued the cyclin D1 mRNA level (Fig. 3F right). We also observed the same phenomenon in ACHN cells (Fig. 3G).

**Figure 3.**
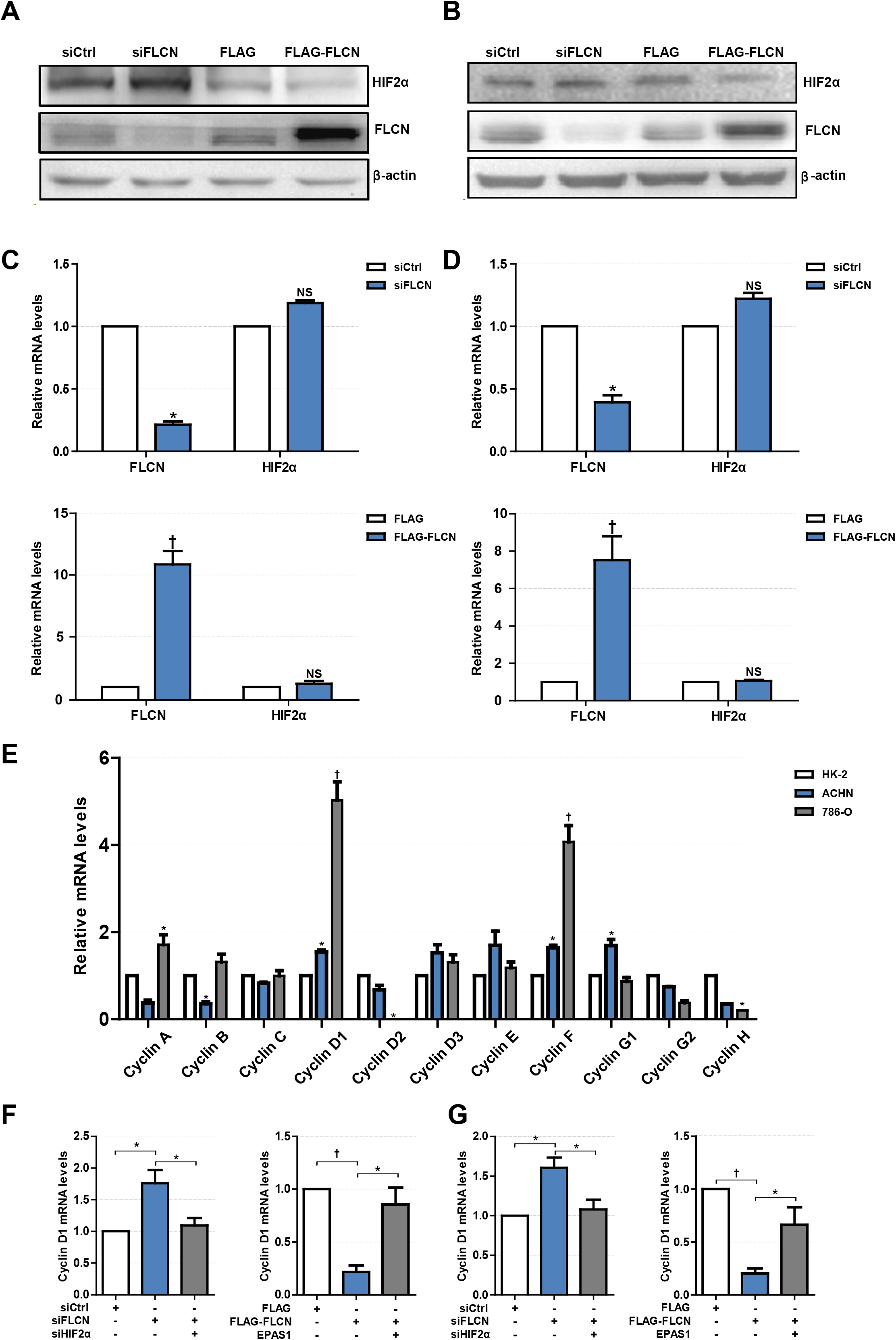
FLCN regulates ccRCC proliferation and cell cycle through HIF2α. (A-D) The regulatory effect of FLCN on HIF2α protein (A,B) and mRNA (C,D) in ccRCC cells was detected by Western blotting and RT-qPCR, respectively. 786-O and ACHN cells were transfected with control siRNA or siFLCN for 48 or 24h.(E) RT-qPCR analysis of cyclin family mRNA expression in ccRCC cell lines. ccRCC cell lines including 786-O and ACHN were used in this experiment. Normal renal tubular epithelium cell line HK-2 was used as control. (F,G) RT-qPCR analysis of cyclin D1 mRNA expression in 786-O (F) and ACHN (G) cell lines. The cells were co-transfected with siFLCN and siHIF2α or FLCN and EPAS1 overexpression plasmid for 24h. The data was from three independently repeated experiments. *: *P*<0.05, †: *P*<0.01.

### 3.4 FLCN knockdown restricts HIF2α degradation of ccRCC cells

In order to confirm whether FLCN bound to HIF2α directly, we further identified the endogenous protein interaction between renal cancer cells and normal renal tubular epithelial cells by immunoprecipitation (Fig. 4A). Whereas 786-O/ACHN cells underwent markable reduction or increase in FLCN, the abundance of HIF2α mRNA was not altered greatly (Figure. 3C and D). Thus, we concluded that instead of transcription-dependent mechanism, FLCN might modulate HIF2α expression by promoting its degradation process. We found that FLCN knockdown inhibited the HIF2α protein level with CHX stimulation in 786-O (Fig. 4B and C top) and ACHN cells (Fig. 4B and C bottom). To explore the causes for the above phenomenon, the cells were treated with KU0063794 (mTOR inhibitor) for different times, and then with or without ubiquitinated proteasome inhibitors to verify whether FLCN degraded HIF2α through the ubiquitination pathway. After treatment with KU0063794, inhibition of mTOR was also shown to retard HIF2α degradation in 786-O (Fig. 4D and E top) and ACHN cells (Fig. 4D and E bottom).

**Figure 4.**
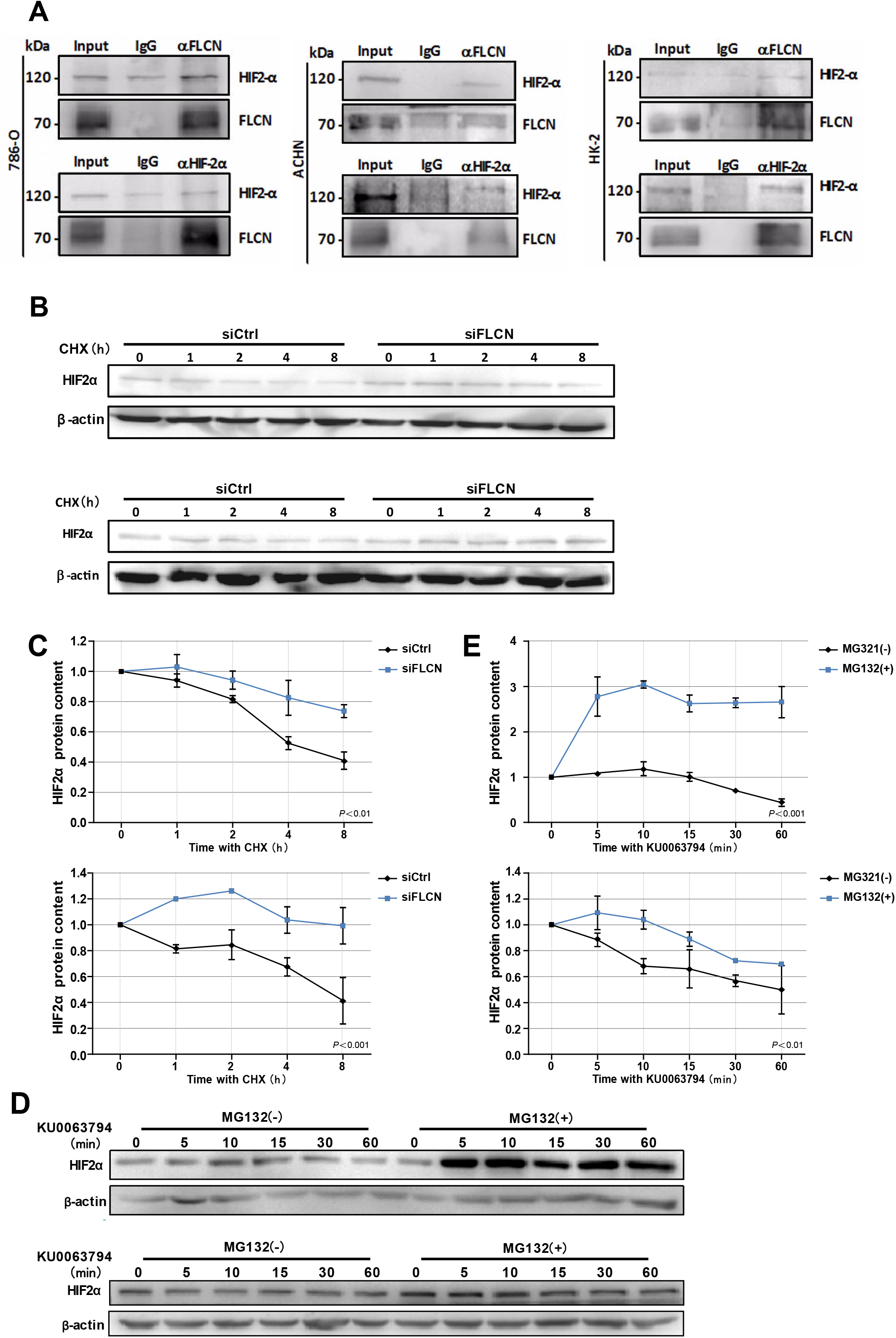
FLCN knockdown restricts HIF2α degradation of ccRCC cells. (A) Western blot showing FLCN co-IP HIF2α or HIF2α co-IP FLCN in 786-O, ACHN and HK-2 cells. IgG as a control. (B,C) 786-O and ACHN cells transfected with control siRNA or siFLCN overnight. In addition to its protein synthesis blocked with cycloheximide (CHX, 10 μg/mL) for the indicated times. Then, the cells were lysed and HIF2α level was determined by Western blotting (B). HIF2α bands were quantified and normalized against β-actin (C). (D,E) 786-O and ACHN cells in addition to its Ubiquitination inhibitors with DMSO or MG-132 (5μg/mL), the cells were then incubated KU0063794 (10 ng/mL) for the indicated times. Then, the cells were lysed and HIF2α level was determined by Western blotting (D). HIF2α bands were quantified and normalized against β-actin (E). The data was from three independently repeated experiments. *: *P*<0.05, †: *P*<0.01.

### 3.5 FLCN knockdown induces HIF2α transportation to nucleus

In view of the above findings, we further explored the influence of FLCN on the function of HIF2α in cell proliferation. Firstly, we examined the exact location of FLCN and HIF2α in the cells, and found that these two molecules were co-localized (Fig. 5A). Then immunofluorescence results showed that knockdown of FLCN promoted HIF2α to enter the nucleus (Fig. 5B). After 46-48 hours (approximately the time of cell division generation) of FLCN knockdown, HIF2α turned into massive in the 786-O cell nucleus at this time point (Fig. 5D) compared to the control group (Fig 5C). Therefore, we verified the validity of this conclusion again by Western blotting in 786-O cells. The results showed visible nucleus HIF2α, which was further increased by FLCN knockdown 46-48h (Fig. 5E bottom), and a slight decrease in expression of HIF2α in the cytoplasma (Fig. 5E top). The same experiment was also repeated in ACHN cells (Fig. 5F), and similar results were obtained.

**Figure 5.**
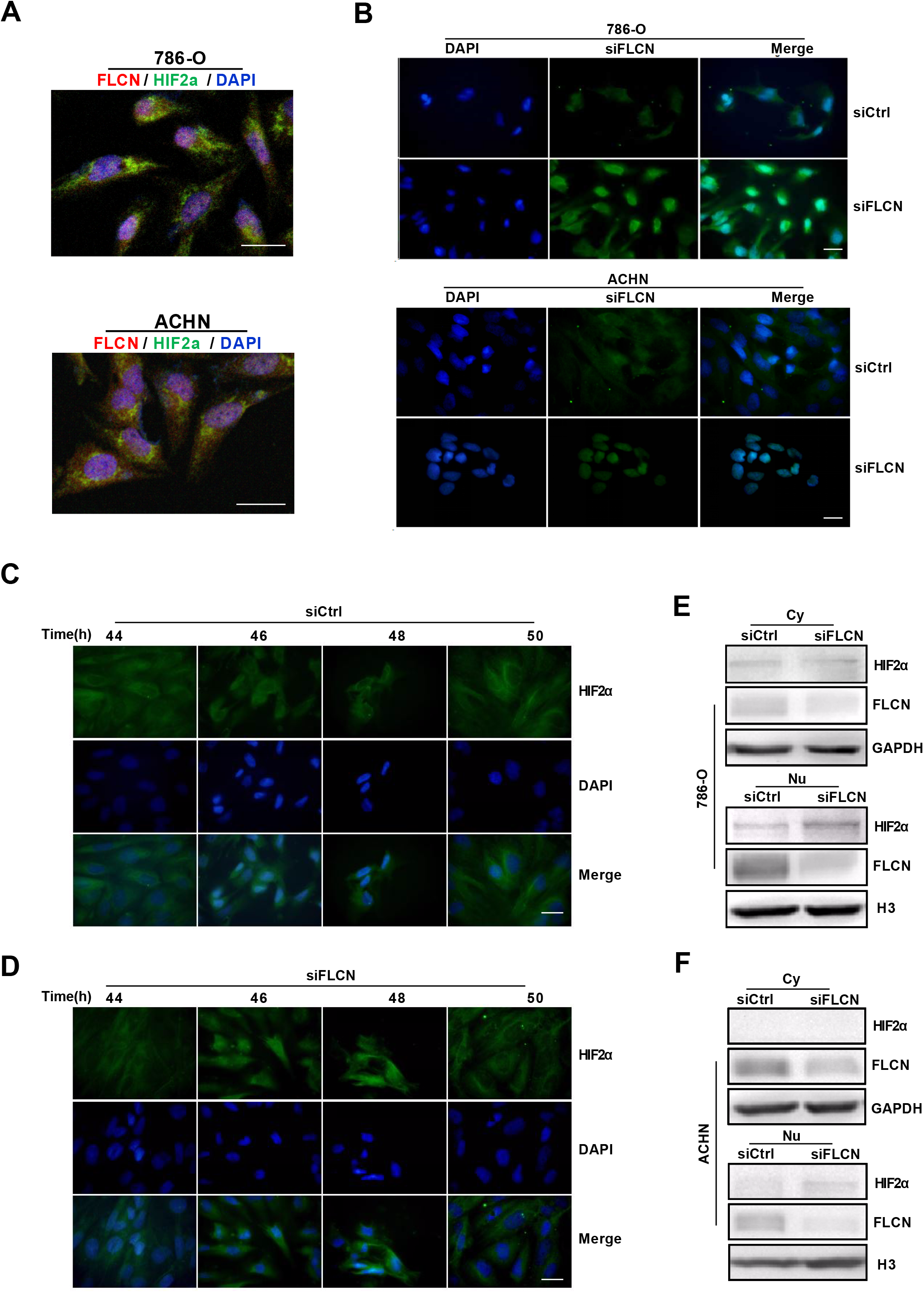
FLCN knockdown induces HIF2α transportation to nucleus. (A) Double immunofluorescence analysis was performed for FLCN and HIF2α. DAPI fluorescence was shown as blue, FLCN immunofluorescence was shown as red, and HIF2α immunofluorescence was shown as green. (B) 786-O and ACHN cells transfected with control siRNA or siFLCN for 46h or 48h. Representative microscopy images of two cells stained immunofluorescence for HIF2α, cell nuclei were labeled with DAPI. (C-F)786-O cells transfected with control siRNA(C) or siFLCN(D) for the indicated times. Then, the cells representative microscopy images of 786-O cells stained immunofluorescence for HIF2α, cell nuclei were labeled with DAPI. The extracts of cytoplasm (E) and nucleus (F) were subjected to Western blotting to detect HIF2α. GAPDH or Histone 3 was as control for cytoplasm or nucleus part. Three independent experiments were carried out. Scale bar, 20μm.

### 3.6 HIF2α mediates FLCN◻induced cell invasion via PI3K/mTORC2 signaling pathways

We examined MMP9 protein and mRNA levels by Western blotting and RT-PCR respectively in 786-O cells transfected with siFLCN or FLCN plasmids (Fig.6A and 6B). We then explored the mechanism of how PI3K/mTORC2 signaling regulated the invasion of FLCN. Recent studies have showed that mTORC is activated in kidney tumors that lack FLCN ^10^. Similarly, Tiffiney R. Hartman et al. reported that downregulation of BHD reduced the TORC1 activity ^21^, which suggested a close relationship between FLCN and mTORC. It was also reported that mTORC promoted HIF expression, mTORC2 focused on HIF2α, and HIF1α was sensitive to rapamycin but HIF2α was tolerant ^22^. Based on the above observations, our study hypothesized that FLCN relied on the PI3K/mTORC signaling pathway to achieve regulation of HIF2α. Firstly, we detected the expression of *FLCN* and found that silencing or overexpression of this gene had no effect on p-AKT S473, an indicator of TORC1 activity (Fig. 6C). Then, we treated 786-0 cells with LY294002, an inhibitor of PI3K, and found that that LY294002 treatment significantly increased HIF2α expression (Fig. 6D). In order to verify the clear upstream and downstream of FLCN, we treated the cells with KU0063794 (mTORC inhibitor), which promoted FLCN expression (Fig. 6E), but could not find significant change in FLCN after adding AKT inhibitor MK2206 (Fig. 6F). These results indicated that mTORC was upstream of FLCN/HIF2α. Finally, we found that rapamycin stimulation did not affect FLCN expression (Fig.6G), which further validated FLCN regulation of HIF2 via mTORC2, but not mTORC1.

**Figure 6.**
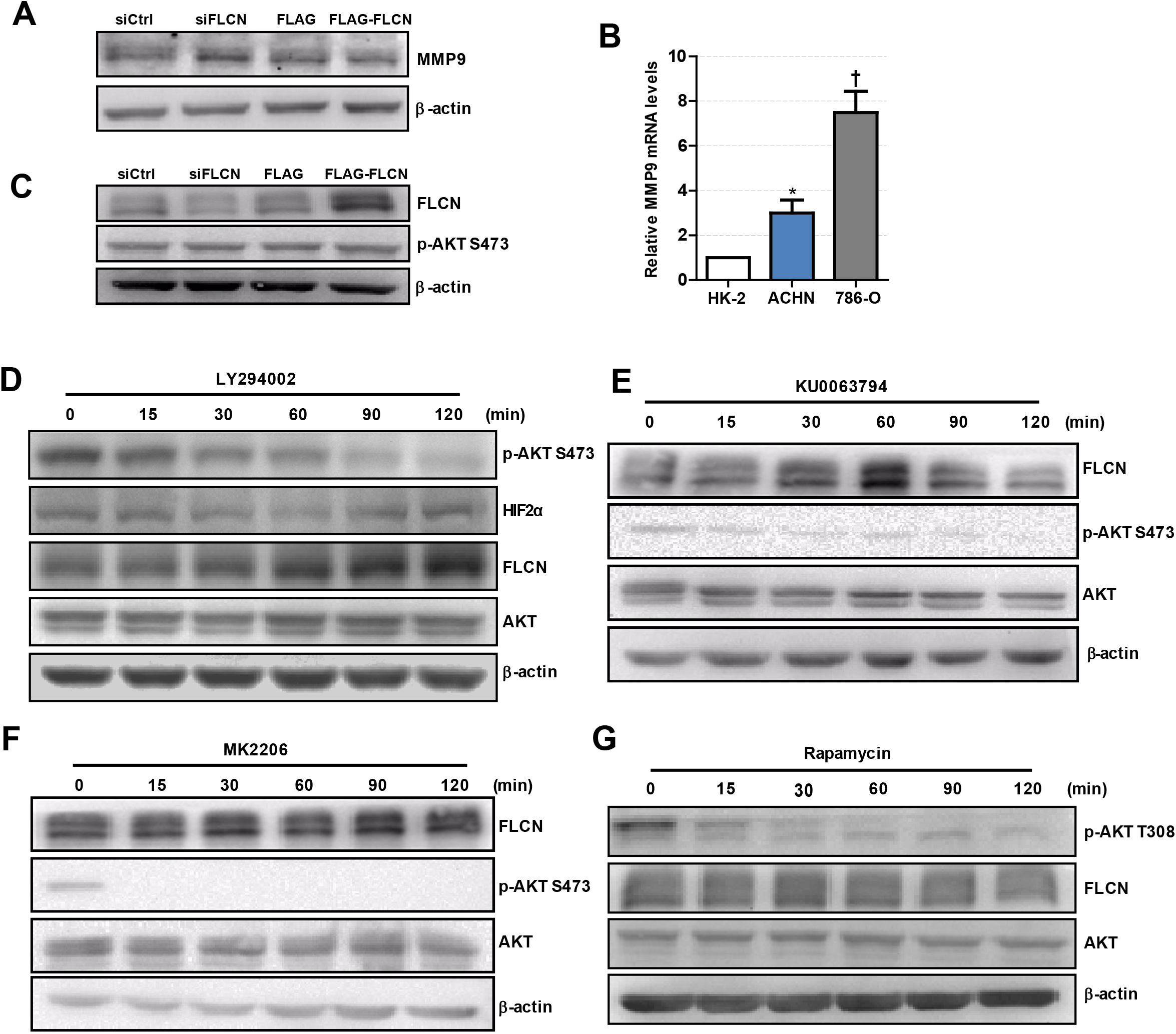
FLCN regulates ccRCC invasion via PI3K/mTORC2/HIF2α signaling pathway. (A) 786-O cells transfected with control siRNA or siFLCN the protein levels of MMP9 were detected by Western blotting and (B) mRNA was detected by RT-qPCR. (C) 786-O cells transfected with control siRNA or siFLCN the protein levels of p-AKT were detected. (D-G) 786-O cells were incubated with LY294002 (PI3K inhibitor), KU0063794 (mTORC inhibitor), MK2206 (AKT inhibitor) and Rapamycin (mTORC1 inhibitor) for 0-120 min, protein levels of FLCN, P◻AKT and AKT were examined. The data was from three independently repeated experiments. The data was repeated from three independent experiments. *: *P*<0.05, †: *P*<0.01.

### 3.7 FLCN is marginally expressed in human renal cancer samples and correlated with HIF2α expression

In order to investigate whether the *in vitro* experimental findings were consistent with the pathogenesis and progression of renal cancer in humans, we examined FLCN and HIF2α expression patterns in renal cancers (75 paired cases) by a tissue microarray (Fig. 7A). Immunohistochemistry results indicated that FLCN was marginally expressed in tumor tissues compared with matched para-cancerous tissues (Figure 7B) but HIF2α was reversed. The immunoreactive score (IRS) was calculated as the intensity of the staining reaction multiplied by the percentage of positive cells. Based on the analysis of 75 groups of renal cancer tissues and para-cancerous tissues, we found that the FLCN expression level was low in tumor tissues, while HIF2α expression level was in a reversed pattern (Figure 7C and 7D). Overall, the clinical data supported our *in vitro* results and revealed a negative link between FLCN and HIF2α protein expression in renal cancer.

**Figure 7.**
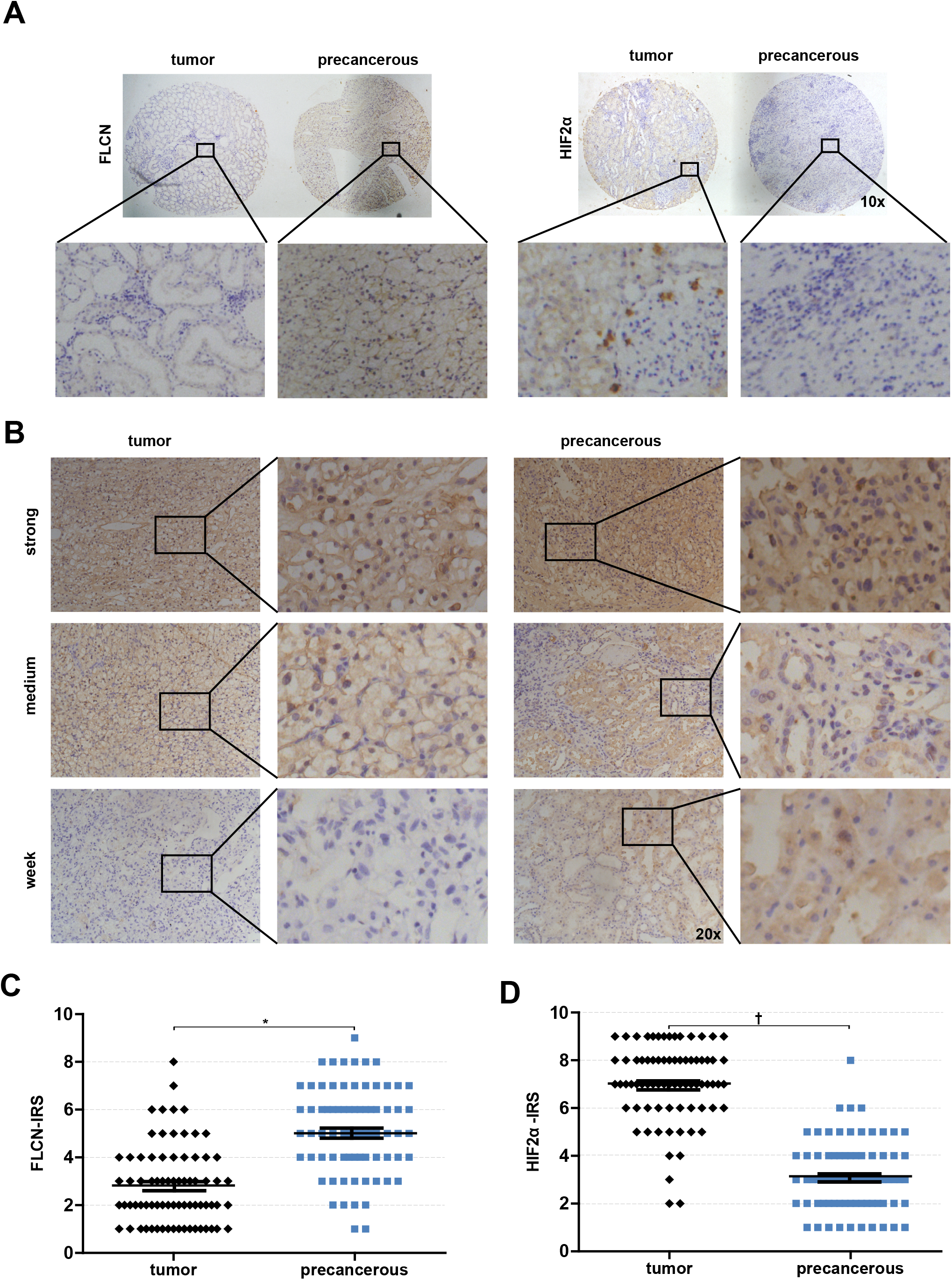
Analysis of FLCN and HIF2α expressions in renal cancer tissues. (A) Representative of malignant renal cancer tissue and paracancerous tissue stained for FLCN and HIF2α are shown. (B) Representative images of FLCN stained in renal cancer tissues are shown. The positive staining of FLCN is shown in brown color and the cell nuclei were counterstained with hematoxylin. (C-D) Analysed for FLCN and HIF2α staining in renal cancer tissue (n = 75) and paracancerous tissue (n = 75) by IRS scores. *: *P*<0.05, †: *P*<0.01.

## 4. Discussion

There are many reports on genes associated with kidney cancer pathogenesis ^23,24^, including *VHL, FH, MET, FLCN,* etc. ^25,26^. The *FLCN* gene is a tumor suppressor ^27^, but the mechanism by which *FLCN* deficiency causes renal cancer is not completely understood. The PI3K/AKT/mTOR pathway, which drives cell proliferation, motility, and migration in a variety of malignancies ^23,28^. Loss of FLCN is linked with activity of the AKT/mTORC pathway ^29,30^, leading to Birt-Hogg-Dubé (BHD) autosomal dominant syndrome, increasing the risk of hybrid oncocytic RCC and pulmonary cysts ^21,31^. It can be seen that these four genes are closely related to HIF and RCC. In summary, we decided to focus on FLCN and study the effects of its inactivation on ccRCC, HIF and the related signaling pathways involved.

A current literature reports that there is a controversy about the relationship between FLCN and HIFα ^4^. HIFα is highly expressed in BHD tumors, and FLCN knockout leads to increased activity of HIFα. However, most of the studies just stayed in the phenomenon. The mechanism is not discussed and the specific relationship between FLCN and HIF1α/HIF2α is unclear. The main points are as follows: RS Preston et al. thought the absence of the FLCN gene did not affect HIF1α levels under normoxia/hypoxia, and it is also mentioned that HIF2α is highly expressed under hypoxic conditions ^32^. In another report, IHC results indicate that FLCN deficiency increases HIF1α and its downstream expression ^33^. A document in the same year claimed that HIF1α increased in the lack of FLCN and promoted tumor growth ^34^. In addition, our study showed that silencing FLCN enhanced the expression of HIF2α protein, and implicated a novel molecular mechanism that FLCN interacted HIF2α and impacted ccRCC physiological activity. In the process, we also detected the mRNA level of HIF2α, and found that there was no significant change, suggesting that FLCN did not regulate HIF2α at the transcriptional level but might control the degradation of HIF2α.

In the experiment, it could be clearly seen that the RNA and protein levels of FLCN in ccRCC were relatively low compared to normal renal tubular epithelial cells. This again proved that *BHD* acted as a tumor suppressor gene, but its role might be more prone to maintain the normal physiological function of the cells. And *BHD* deletion activated the downstream signaling pathway, resulting in tumor formation, rather than saving the deterioration of existing tumor cells. Based on the above observations, we transfected siFLCN with ccRCC to demonstrate that FLCN deficiency had a detailed mechanism for inducing renal cell carcinoma. At the same time, FLCN overexpression experiments were also performed to verify whether the increasing FLCN expression levels in renal tumors caused by other factors contributed to the alleviation of tumor cell deterioration. In the experiment, we selected 786-O and ACHN two ccRCC cell lines, of which 786-O was a cell line with VHL gene deletion, and HIF2α was still highly expressed under normoxia, which was more convenient for us to observe the results. Then, we added CHX interfering protein translation to the cell line knocked down by FLCN. The results showed that compared with the control group, the content of HIF2α in the FLCN interference group could be maintained at a relatively stable level within 0-8 hours after the addition of CHX, indicating that FLCN might accelerate the degradation of HIF2α protein. We hypothesized that this mechanism might have some similarities with VHL ubiquitination and degradation of HIF1α ^35–37^. To further verify the specific mechanism of this degradation, we added the mTORC inhibitor to up-regulate the FLCN, and at the same time added the proteasome inhibitor MG-132 to inhibit the ubiquitination degradation pathway. The results showed that although increased FLCN could reduce HIF2α, but after adding MG132 the expression of HIF2α was significantly increased, which indicated that FLCN was likely to achieve the degradation of HIF2α by ubiquitination, thereby inhibiting the proliferation of tumors.

In addition, during the experiment, we also unexpectedly found that the transcription factor HIF2α in the siFLCN group increased significantly in the nucleus at 46-48 hours compared with the control group. In order to study the occurrence of this phenomenon, it was caused by the FLCN knockdown which promoted the acceleration of HIF2α into the nucleus, rather than the increase of the total protein of HIF2α. We transfected siFLCN in ccRCC and observed the cell morphological phenomenon from 0 h with immunofluorescence, and then detected the distribution of HIF2α in the cells every two hours (only the distribution of 44 h-50 h was shown in the figures). The results showed that HIF2α in the siFLCN group of the 786-O cell line began to enter the nucleus in 46-48 hours, rather than gradually increasing with time. At the same time, the distribution and content of HIF2α in the nucleus of the control group did not change significantly. This indicated that FLCN could not only bound and degraded HIF2α, but also participated in the regulation of HIF2α nuclear import: silencing FLCN, HIF2α might move into the nucleus and activated downstream related tumor signaling pathways.

In conclusion, we demonstrate the specific mechanism by which FLCN regulates HIF2α-induced renal cancer cells proliferation, migration, and invasion. FLCN interacts with HIF2α and promotes HIF2α degradation in human renal cancer cells. Although the current study has contributed to the mechanistic understanding the role of FLCN regulates renal cancer cell physiological activities, the issue as to how FLCN precisely adjusts HIF2 degradation in ccRCC cells is unlikely to be settled in this paper. Our findings point out that FLCN/HIF2α expression may be a novel therapeutic target for preventing renal cancer proliferation and invasion.

## Abbreviations

FLCN: folliculin
VHL: Von Hippel - Lindau
MMP9: matrix metalloproteinase 9
CHX: cycloheximide
ccRCC: clear cell renal cell carcinoma

## 5. Acknowledgements

This work was supported by the National Natural Science Foundation of China to Luo GU (No. 81372319), Yujie Zhang (No. 81602561) and Jun Du (No. 81773107), and a Project Funded by Jiangsu Collaborative Innovation Center for Cancer Personalized Medicine to Luo Gu.

## 6. Conflict of interest

The authors confirm that there are no conflicts of interest.

## References

1. Kovacs G, Akhtar M, Beckwith BJ, et al. The Heidelberg classification of renal cell tumours. The Journal of Pathology. 1997;183(2):131–133.

2. Linehan WM, Srinivasan R, Schmidt LS. The genetic basis of kidney cancer: a metabolic disease. Nat Rev Urol. 2010;7(5):277–285.

3. Toro JR, Wei MH, Glenn GM, et al. BHD mutations, clinical and molecular genetic investigations of Birt-Hogg-Dube syndrome: a new series of 50 families and a review of published reports. J Med Genet. 2008;45(6):321–331.

4. Hasumi Y, Baba M, Ajima R, et al. Homozygous loss of BHD causes early embryonic lethality and kidney tumor development with activation of mTORC1 and mTORC2. Proc Natl Acad Sci U S A. 2009;106(44):18722–18727.

5. Khoo SK, Bradley M, Wong FK, Hedblad MA, Nordenskjold M, Teh BT. Birt-Hogg-Dube syndrome: mapping of a novel hereditary neoplasia gene to chromosome 17p12-q11.2. Oncogene. 2001;20(37):5239–5242.

6. Schmidt LS, Linehan WM. Molecular genetics and clinical features of Birt-Hogg-Dube syndrome. Nat Rev Urol. 2015;12(10):558–569.

7. Schmidt LS, Warren MB, Nickerson ML, et al. Birt-Hogg-Dube syndrome, a genodermatosis associated with spontaneous pneumothorax and kidney neoplasia, maps to chromosome 17p11.2. Am J Hum Genet. 2001;69(4):876–882.

8. Baba M, Furihata M, Hong SB, et al. Kidney-targeted Birt-Hogg-Dube gene inactivation in a mouse model: Erk1/2 and Akt-mTOR activation, cell hyperproliferation, and polycystic kidneys. J Natl Cancer Inst. 2008;100(2):140–154.

9. Hudon V, Sabourin S, Dydensborg AB, et al. Renal tumour suppressor function of the Birt-Hogg-Dube syndrome gene product folliculin. J Med Genet. 2010;47(3):182–189.

10. Chen J, Futami K, Petillo D, et al. Deficiency of FLCN in mouse kidney led to development of polycystic kidneys and renal neoplasia. PLoS One. 2008;3(10):e3581.

11. Hartman TR, Nicolas E, Klein-Szanto A, et al. The role of the Birt-Hogg-Dube protein in mTOR activation and renal tumorigenesis. Oncogene. 2009;28(13):1594–1604.

12. Schmidt LS, Linehan WM. Clinical Features, Genetics and Potential Therapeutic Approaches for Birt-Hogg-Dube Syndrome. Expert Opin Orphan Drugs. 2015;3(1):15–29.

13. Gallou C, Joly D, Mejean A, et al. Mutations of the VHL gene in sporadic renal cell carcinoma: definition of a risk factor for VHL patients to develop an RCC. Hum Mutat. 1999;13(6):464–475.

14. Kondo K, Kaelin WG, Jr. The von Hippel-Lindau tumor suppressor gene. Exp Cell Res. 2001;264(1):117–125.

15. Aleksic T, Browning L, Woodward M, et al. Durable Response of Spinal Chordoma to Combined Inhibition of IGF-1R and EGFR. Front Oncol. 2016;6:98.

16. Du W, Zhang L, Brett-Morris A, et al. HIF drives lipid deposition and cancer in ccRCC via repression of fatty acid metabolism. Nat Commun. 2017;8(1):1769.

17. Maranchie JK, Zhan Y. Nox4 is critical for hypoxia-inducible factor 2-alpha transcriptional activity in von Hippel-Lindau-deficient renal cell carcinoma. Cancer Res. 2005;65(20):9190–9193.

18. Pawlus MR, Wang L, Ware K, Hu CJ. Upstream stimulatory factor 2 and hypoxia-inducible factor 2alpha (HIF2alpha) cooperatively activate HIF2 target genes during hypoxia. Mol Cell Biol. 2012;32(22):4595–4610.

19. Zhu Y, Shen T, Liu J, et al. Rab35 is required for Wnt5a/Dvl2-induced Rac1 activation and cell migration in MCF-7 breast cancer cells. Cell Signal. 2013;25(5):1075–1085.

20. Zhang Y, Du J, Zheng J, et al. EGF-reduced Wnt5a transcription induces epithelial-mesenchymal transition via Arf6-ERK signaling in gastric cancer cells. Oncotarget. 2015;6(9):7244–7261.

21. Bonora M, Wieckowsk MR, Chinopoulos C, et al. Molecular mechanisms of cell death: central implication of ATP synthase in mitochondrial permeability transition. Oncogene. 2015;34(12):1608.

22. Toschi A, Lee E, Gadir N, Ohh M, Foster DA. Differential dependence of hypoxia-inducible factors 1 alpha and 2 alpha on mTORC1 and mTORC2. J Biol Chem. 2008;283(50):34495–34499.

23. Blume-Jensen P, Hunter T. Oncogenic kinase signalling. Nature. 2001;411(6835):355–365.

24. Zbar B, Klausner R, Linehan WM. Studying cancer families to identify kidney cancer genes. Annu Rev Med. 2003;54:217–233.

25. Davis CF, Ricketts CJ, Wang M, et al. The somatic genomic landscape of chromophobe renal cell carcinoma. Cancer Cell. 2014;26(3):319–330.

26. Hsieh JJ, Purdue MP, Signoretti S, et al. Renal cell carcinoma. Nat Rev Dis Primers. 2017;3:17009.

27. Nickerson ML, Warren MB, Toro JR, et al. Mutations in a novel gene lead to kidney tumors, lung wall defects, and benign tumors of the hair follicle in patients with the Birt-Hogg-Dube syndrome. Cancer Cell. 2002;2(2):157–164.

28. Vivanco I, Sawyers CL. The phosphatidylinositol 3-Kinase AKT pathway in human cancer. Nat Rev Cancer. 2002;2(7):489–501.

29. Cash TP, Gruber JJ, Hartman TR, Henske EP, Simon MC. Loss of the Birt-Hogg-Dube tumor suppressor results in apoptotic resistance due to aberrant TGFbeta-mediated transcription. Oncogene. 2011;30(22):2534–2546.

30. Schmidt LS, Linehan WM. FLCN: The causative gene for Birt-Hogg-Dube syndrome. Gene. 2018;640:28–42.

31. Kapitsinou PP, Haase VH. The VHL tumor suppressor and HIF: insights from genetic studies in mice. Cell Death Differ. 2008;15(4):650–659.

32. Preston RS, Philp A, Claessens T, et al. Absence of the Birt-Hogg-Dube gene product is associated with increased hypoxia-inducible factor transcriptional activity and a loss of metabolic flexibility. Oncogene. 2011;30(10):1159–1173.

33. Hasumi Y, Baba M, Hasumi H, et al. Folliculin (Flcn) inactivation leads to murine cardiac hypertrophy through mTORC1 deregulation. Hum Mol Genet. 2014;23(21):5706–5719.

34. Khabibullin D, Medvetz DA, Pinilla M, et al. Folliculin regulates cell-cell adhesion, AMPK, and mTORC1 in a cell-type-specific manner in lung-derived cells. Physiol Rep. 2014;2(8).

35. Epstein ACR, Gleadle JM, McNeill LA, et al. C. elegans EGL-9 and Mammalian Homologs Define a Family of Dioxygenases that Regulate HIF by Prolyl Hydroxylation. Cell. 2001;107(1):43–54.

36. Ivan M, Kondo K, Yang H, et al. HIFalpha targeted for VHL-mediated destruction by proline hydroxylation: implications for O2 sensing. Science. 2001;292(5516):464–468.

37. Jaakkola P, Mole DR, Tian YM, et al. Targeting of HIF-alpha to the von Hippel-Lindau ubiquitylation complex by O2-regulated prolyl hydroxylation. Science. 2001;292(5516):468–472.

